# GLARE: Discovering Hidden Patterns in Spaceflight Transcriptome Using Representation Learning

**DOI:** 10.1101/2024.06.04.597470

**Authors:** DongHyeon Seo, Hunter F. Strickland, Mingqi Zhou, Richard Barker, Robert J Ferl, Anna-Lisa Paul, Simon Gilroy

## Abstract

Spaceflight studies present novel insights into biological processes through exposure to stressors outside the evolutionary path of terrestrial organisms. Despite limited access to space environments, numerous transcriptomic datasets from spaceflight experiments are now available through NASA’s GeneLab data repository, which allows public access to these datasets, encouraging further analysis. While various computational pipelines and methods have been used to process these transcriptomic datasets, learning-model-driven analyses have yet to be applied to a broad array of such spaceflight-related datasets. In this study, we propose an open-source pipeline, GLARE: GeneLAb Representation learning pipelinE, which consists of training different representation learning approaches from manifold learning to self-supervised learning that enhance the performance of downstream analytical tasks. We illustrate the utility of GLARE by applying it to gene-level transcriptional values from the results of the CARA spaceflight experiment, an Arabidopsis root tip transcriptome dataset that spanned light, dark, and microgravity treatments. We show that GLARE not only substantiated the findings of the original study concerning cell wall remodeling but also revealed additional patterns of gene expression affected by the treatments, including evidence of hypoxia. This work suggests there is great potential to supplement the insights drawn from initial studies on spaceflight omics-level data through further machine-learning-enabled analyses.

## 1 Introduction

Spaceflight studies present unprecedented insights into biological processes through exposure to unique environmental stressors that have not been experienced by any form of life on Earth. In response to the spaceflight environment, organisms initiate specific transcriptional responses to novel conditions. Thus, one key to understanding how biology responds to spaceflight stressors like microgravity, radiation, and hypoxia is through transcriptomic analysis to study the gene expression profiles that drive physiological adaptation triggered by the spaceflight environment (Mustroph et al., 2010). Space-related transcriptional studies have now also broadened into multi-omic spaceflight investigations that are well-suited to multiple rounds of analysis facilitated by the publicly available datasets in the NASA GeneLab database.

The importance of studying plant biology specifically in space has been identified both for exploring the fundamental responses of biology to the spaceflight environment and at a very practical level for developing bio-regenerative life support systems for long-term space exploration (Rutter et al., 2020; Fu et al., 2016). Understanding of transcriptomic and physiological changes elicited in plants by spaceflight conditions through analyzing transcriptional and other –omic patterns is therefore a focus of much current plant space biology experimentation (e.g., Paul et al. (2013); Villacampa et al. (2021)). For example, the CARA (Characterizing Arabidopsis Root Attraction) experiment was designed to compare the spaceflight transcriptome responses between different genotypes of *Arabidopsis thaliana’s* root tips under various conditions (Paul et al., 2017). This experiment explored the patterns of gene expression from root tip cells in the spaceflight environment on the International Space Station (ISS) with comparable ground controls, and the lighting sub-environments among three different genotypes. While these kinds of experiments in plant space biology have provided many key insights, they have so far largely relied upon the primary transcriptomic analysis of the original research team. To provide a pipeline that can be applied to increase the depth of transcriptomic analyses for previous and future spaceflight experiments, we introduce GLARE: GeneLab Representation learning pipelinE. This pipeline consists of multiple methods of data visualization and projection, representational learning, and post hoc analyses, provided in a pre-built series of codes targeted for use with datasets from the Genelab Data System (GLDS). We show the utility of the GLARE by applying it to the CARA dataset to illustrate how applying novel machine-learning methods to transcriptomic datasets extends insights beyond the original transcriptomic analysis of this data.

Our analysis pipeline applies state-of-the-art representation learning models to find underlying patterns in the FPKM values (fragments per kilobase of transcript per million mapped fragments) that are proportional to the abundance of each loci’s transcript. The application of representation learning models enhances the characterization of data points, thereby improving downstream tasks such as clustering. This approach facilitates the grouping of genes with similar attributes, offering the possibility to reveal new and significant biological insights (Eisen et al., 1998) and providing a robust foundation for analytical methodologies, such as further investigation of the effects of spaceflight on, e.g., phytohormone signaling and associated physiological phenotypes (Abts et al., 2017; Ferl and Paul, 2016; Iqbal et al., 2017). Moreover, considering that the CARA experiment also utilized lighting sub-environments, we can shine further light on the potential spaceflight effects and interactions with this factor that may have been overlooked in past studies. Overall, the GLARE method will provide insights to better understand plant behavior in the spaceflight environment based on its endogenous and exogenous cues.

## 2 Materials and Methods

### 2.1 GeneLab Data System and Data Entries

The Genelab Data System (GLDS) is a public, space-related -omics data repository, which curates data from a wide variety of species and experimental spaceflight conditions (Ray et al., 2018). GLDS obtains spaceflight-related –omics datasets from multiple locations such as the Gene Expression Omnibus (GEO), European Bioinformatics Institute (EBI), publications directly, and others (Ray et al., 2018). This data is then cataloged with the relevant metadata, such as protocols, payload numbers, and experimental variables, and made available as an Open Science Dataset (OSD) in NASA’s Open Science Data Repository (OSDR). The CARA dataset (OSD-120; https://osdr.nasa.gov/bio/repo/data/studies/OSD-120) was chosen from the GLDS to use as a test case for analysis using GLARE due to its many experimental conditions. The CARA experiments were conducted with three ecotypes/genotypes of *Arabidopsis thaliana*: wild-type Wassilewskija (WS), wild-type Columbia-0 (Col-0), and a mutant in the *PHYTOCHROME D* gene in the Col-0 background (*PHYD*) (Paul et al., 2017). Briefly, these genotypes were planted on gel media in Petri dishes and grown in either ambient light conditions or in the dark on the ISS for 11 days; Parallel controls were performed on the ground. After the 11 days, germinated seedlings were photographed and collected into Kennedy Space Center Fixation Tubes (KFTs; Ferl et al. (2011)) containing RNAlater. Seedlings preserved in RNAlater were returned to Earth frozen, and then the roots were dissected into the last 2 mm of the tip for the light-grown plants and the last 1 mm for the dark-grown plants. RNA was extracted and sent to the Interdisciplinary Center for Biotechnology Research (ICBR), University of Florida, for RNA sequencing using a NextSeq 500 system, producing ∼40 million paired-end reads per sample. Finally, these paired-end reads were mapped to the TAIR10 *A. thaliana* reference genome using Spliced Transcripts Alignment to a Reference (STAR) software, and differential expression was performed using the Cufflinks tool (Dobin et al., 2013; Trapnell et al., 2012).

### 2.2 High-dimensional Data Analysis

**Overview**: Statistical methods have been widely integrated into the bioinformatics pipelines in multi-omics studies for analyzing the data. Specifically, due to multi-omics datasets having complex data topology, dimension reduction and clustering are two commonly used techniques for further investigation (Rappoport and Shamir, 2018). GLARE capitalizes upon such approaches. For example, Principal Component Analysis (PCA) and Factor Analysis are methods with widespread application for dimensionality reduction (Zeng and Lumley, 2018). After achieving a statistical representation of the dataset with these dimensionality reduction techniques, clustering methods are utilized to group similar representations to uncover underlying patterns within the dataset. Among these, K-means and hierarchical clustering are featured as two of the most favored methodologies (Hulot et al., 2020).

#### 2.2.1 Learning Data Representations

While PCA is popularly used for its simplicity and efficiency, it has its limits for losing essential non-linear features through linear embedding, which often degrades the clustering quality (Gan et al., 2020). Several alternative methods that do not only rely on data point distribution but also leverage latent data structures via learned representations have shown advantages in handling biological data, thereby enhancing clustering precision (Karim et al., 2021). GLARE also uses these approaches in its analyses. These alternatives to PCA include t-distributed Stochastic Neighbour Embedding (t-SNE), a non-linear dimensionality reduction technique particularly adept at preserving local structures within high-dimensional data, and Uniform Manifold APproximation (UMAP) (Van der Maaten and Hinton, 2008; McInnes et al., 2018), a manifold learning approach that efficiently captures complex relationships within the data. However, alternative deep-learning-based approaches for obtaining data representations have been largely neglected in the field of plant space biology, despite the advantage of their ability to capture contextual information from the non-linear mappings. Specifically, this approach of capturing contextual information through complex, higher-level features is known as representation learning or feature extraction (Aljalbout et al., 2018). Therefore, along with PCA, t-SNE, and UMAP, we have investigated the application of Sparse Autoencoder (SAE) as one of the representation learning methods in the GLARE. SAE is an unsupervised learning algorithm based on a neural network that aims to learn an approximation of the identity function that represents the data. The model is trained by encoding the data from its feedforward phase but with sparsity constraints that trigger neurons with activation above a threshold, i.e., those with the largest activation, allowing the discovery of the unique structure in the data (Makhzani and Frey, 2013). While autoencoders are more commonly used for reconstructing the original input data, prior studies show autoencoder as a representation learning approach that works favorably in the context of multi-omics datasets (Chaudhary et al., 2018).

Upon employing multiple approaches to obtain data representation, evaluating these representations is critical to understanding the strengths and limitations of various data representation techniques. Prior research has used several evaluation techniques to assess the fidelity between data representations and the original dataset and the quality of the data representation structure, so we have used these methods in the development of the GLARE approach. Trustworthiness score measures the preservation of local topological structure in the data and is widely applied to evaluate the fidelity and faithfulness of the learned representation, testing its ability to maintain local structure and inherent relationships (Hinton and Salakhutdinov, 2006; Venna and Kaski, 2001). On the other hand, evaluating the efficacy of these data representations in potential downstream tasks, such as classification or clustering, provides crucial insights into their practical utility and generalizability. K-Nearest Neighbors (KNN) classifier can be utilized to test the quality and separability of the data structure in reduced dimension by assessing the class-discriminative information (Van Der Maaten et al., 2009). Furthermore, the Silhouette score is widely used to check the insights into clustering performance and compactness of the data representation (Rousseeuw, 1987). These metrics would help determine when it’s appropriate to use the data representations for specific downstream tasks.

#### 2.2.2 Clustering Data Representations

Within the clustering paradigm, several alternative methods to K-means exist for the effective organization of these representations, and we have explored their application as part of the GLARE. Among these, Gaussian Mixture Models (GMM) with the Expectation-Maximization (EM) algorithm offer a probabilistic framework, wherein each cluster is represented by a Gaussian distribution, facilitating more nuanced cluster assignments (Reynolds et al., 2009). Density-based clustering methods have gained attention with respect to their ability to detect clusters of arbitrary shapes and sizes, thus overcoming some of the limitations associated with distance-based methods (Ester et al., 1996). Notably, an extension of this approach, Hierarchical Density-Based Spatial Clustering of Applications with Noise (HDBSCAN), utilizes a hierarchical approach to density-based clustering to robustly identify clusters at multiple levels with varying densities (Campello et al., 2013). Additionally, spectral clustering presents an alternative approach, leveraging the eigenstructure of the similarity matrix to partition the data into clusters, thereby offering an effective means of characterizing complex structures within the dataset (Ng et al., 2001).

Ensemble clustering is an additional powerful technique that combines these multiple clustering solutions to obtain consensus clusters that are more robust and accurate. Several ensemble clustering methods have been proposed, including Evidence Accumulation Clustering (EAC) (Fred and Jain, 2005), which accumulates evidence from different base clustering algorithms to build a co-association matrix. Applying hierarchical clustering to this matrix derives a final consensus clustering result. Other notable examples include HyperGraph-Partitioning Algorithm (HGPA) (Strehl and Ghosh, 2002), which derives consensus clustering through a partitioning hypergraph where each base clustering set is a hyperedge in a hypergraph, with vertices representing data points. These ensemble techniques have demonstrated their utility in various domains, such as bioinformatics, text mining, and computer vision, where data is often high-dimensional, noisy, and complex (Vega-Pons and Ruiz-Shulcloper, 2011). Therefore, they are strong candidates for equivalent analyses of the often highly complex structures that make up plant transcriptomics datasets. Hence, these approaches are incorporated into the GLARE.

## 3 Results

### 3.1 GLARE: GeneLAb Representation learning pipelinE

We introduce GLARE, a representation learning pipeline designed to empower researchers to move beyond conventional dimensionality reduction techniques in their omics-focused research, such as reliance on PCA or tSNE. GLARE enables the extraction of data representations using a trained learning-based model, thereby allowing the exploration of latent structures to unveil the hidden patterns inherent within the dataset. We first report a verification study by training a classification model on the CARA study’s spaceflight and ground control data, followed by the full analytical pipeline to highlight GLARE’s ability to both confirm patterns revealed in the published primary analyses and reveal novel patterns within the data.

#### 3.1.1 Verification study

Prior to applying the full end-to-end pipeline of GLARE, we implemented a verification study by performing prediction tasks on restructured data. The verification study serves dual purposes: 1) As a validity check for the approach prior to deploying the full GLARE. If the data did not exhibit any learnable and distinctive patterns between the experiment setting that we wanted to compare against, as revealed by making poor predictions on the test set in this verification study, then applying unsupervised methods on that data of interest would be ineffective as the extracted representations would not capture meaningful latent information. 2) The prediction task from the verification study can serve as the foundation for post-pipeline analysis, enabling the incorporation of feature importance explanation schemes, such as SHapley Additive exPlanations (SHAP) (Lundberg and Lee, 2017). The feature importance values can reveal, e.g., within CARA, which genotypes and light conditions contributed the most to the predictions overall, as well as provide more insights into specific genes of interest. Combining these insights with the clustering results from the GLARE should substantially empower researchers in general to see new patterns in their omics-level data.

We focused this verification study on the CARA dataset (OSD-120) to ensure that learnable patterns indeed exist within this spaceflight transcriptome dataset before using it with the GLARE. To enable this analysis, we restructured the original OSD-120 dataset from a wide format to a long format, a process often called ‘data melting’ (Wickham, 2007). This transformation involves reorganizing the unlabeled data so that each experimental factor (such as genotype, spaceflight versus ground control, and lighting regime) becomes a separate categorical variable, or a label, instead of embedding them in the column heading. We, therefore, extracted feature vectors representing each experimental condition and restructured the numerical FPKM data to be indexed by both gene locus and individual experimental factor labels rather than just by gene locus (Figure 1(a)). Essentially, restructured data, *X*_*melted*_, becomes organized to facilitate analysis across multiple experimental dimensions while containing the same information as the original data, *X*, which has 36 numerical features as it contains experiment results from two locations (spaceflight versus ground), under two different light conditions (light versus dark), and for three genotypes (Ws, Col-0, and *PHYD* mutants), with three replicate samples of each.

**Figure 1.**
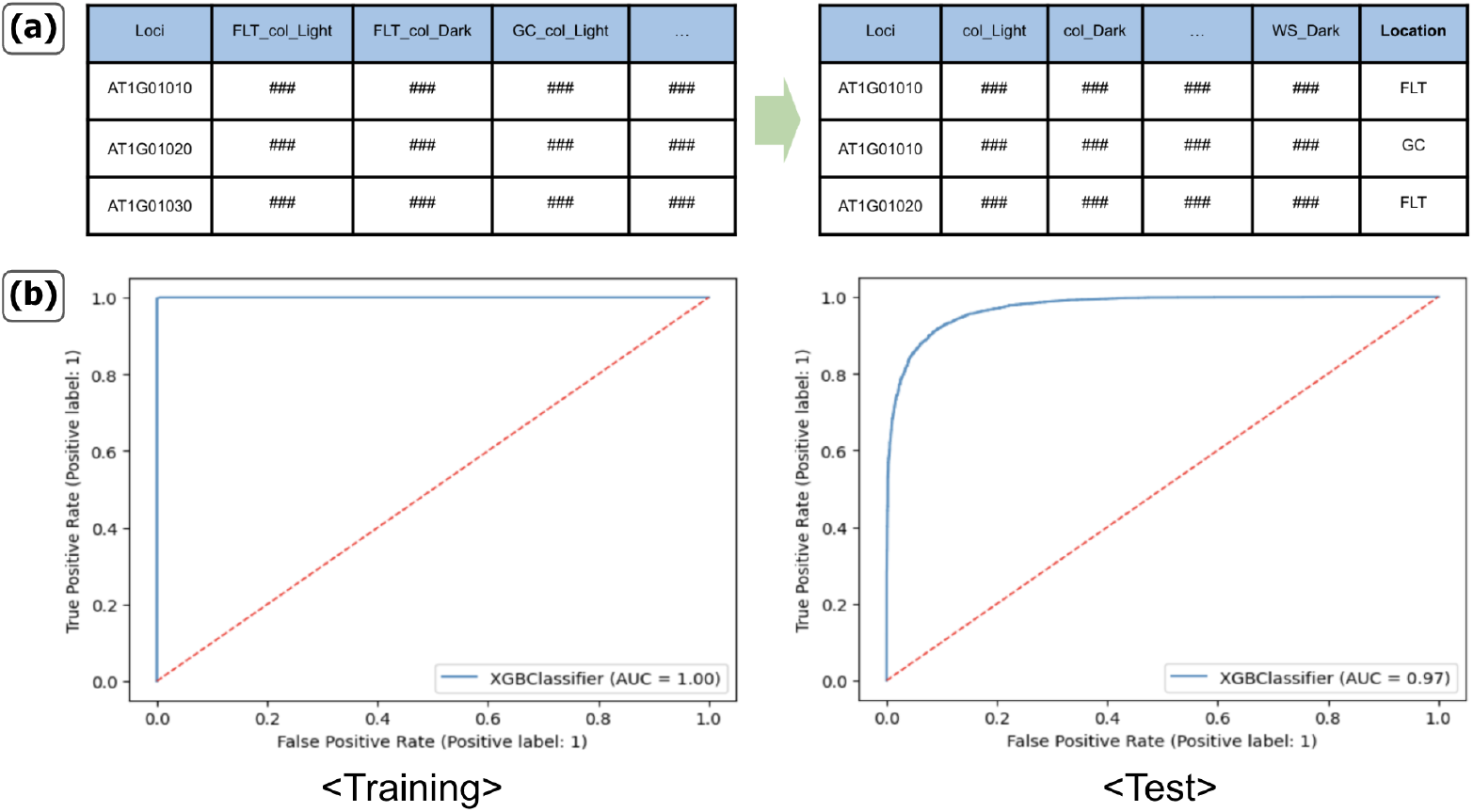
Data melting for data restructuring and supervised learning prediction performances. (a) Illustration showing the restructuring process of our base data model where we make the experiment location a categorical variable. Raw FPKM numerical data (denoted as ###) as reads per locus (e.g., AT1G01010, AT1G01020 across all ∼ 25,000 genes in the Arabidopsis genome) for each experimental sample (e.g., Flight sample of Columbia ecotype grown in the light, FLT_col_Light, or Ground control sample of Columbia ecotype grown in the light, GC_col_Light) is shown left. After restructuring (right table), each gene has two instances, one from the flight and one from the ground sample separately. (b) ROC curves using the training and test dataset on the best-performing data model, which is the base model. Blue line represents the XGBoost classifier, showing the ratio of true positives to false positives in the model predictions from the training data (left) and the test set (right). Red line is the random chance baseline. FLT, spaceflight; GC, ground control.

This restructuring allows the experiment environment to be explicitly indicated through the labels (Xu et al., 2013). We then trained the classification model using XGBoost (Chen and Guestrin, 2016), on the restructured data *X*_*melted*_ using the concatenated feature vectors as the input matrix and the new labels that represent the experiment environment as target labels. High predictive performance, such as accurately assigning a sample to flight versus ground control, would indicate that learnable patterns are indeed present in the original data, thus motivating the use of unsupervised representation learning techniques from GLARE. Figure 1 shows an illustration of how we restructured our dataset through data melting and the prediction performance of the best-performing data model. We compared prediction performances across multiple data melting models on a held-out test set of *X*_*melted*_, which is presented in Table 1. This test set is a random subset that was set aside during training to be used for evaluating the model performance on unseen data. Data models that were tested include our ‘base model’, where we have location labels indicating if the experiments were performed in space (ISS) or on the ground (KSC), having 18 continuous features and one flight versus ground label. We also performed data melting using other experiment settings such as ‘Genotype’ and ‘Light condition’, adding these additional labels to act as further categorical predictors. We found that our base data model without other additional categorical variables yields the highest performance with ∼91% test accuracy on predicting if the experiments were done in space or the ground based on the normalized counts of FPKM values.

**Table 1:** Classification performances on held-out test set using XGBoost on data from different data models (with ± standard deviation). F1-score: The harmonic mean of precision (avoidance of false positives) and recall (avoidance of false negatives), ROC-AUC: Area under the Receiver Operating Characteristic curve, summarizing true positive vs. false positive trade-off. Test Accuracy is the % correctness of predictions for classifying a sample as spaceflight or ground control in the test set using each data model.

| Data Model | Test Accuracy ↑ | F1-score ↑ | ROC-AUC ↑ |
| --- | --- | --- | --- |
| Base data melting model | **$91.29 \pm 0.26$** | **$0.913 \pm 0.002$** | **$0.975 \pm 0.001$** |
| + Genotype data melting | $77.09 \pm 0.14$ | $0.770 \pm 0.002$ | $0.865 \pm 0.001$ |
| + Light data melting | $83.18 \pm 0.07$ | $0.831 \pm 0.001$ | $0.919 \pm 0.001$ |
| + Genotype & Light data melting | $68.69 \pm 0.14$ | $0.686 \pm 0.001$ | $0.772 \pm 0.001$ |

Encouraged by the outcome of this verification study that a machine learning approach should be able to extract potentially novel features from spaceflight datasets, we applied GLARE. Prior to the full pipeline, we implemented a cluster-based outlier detection method (detailed in Supplementary Figure S1) to enhance the data quality. This preprocessing step identified and eliminated three anomalous genes (AT1G0759, AT3G41768, ATMG00020), thereby ensuring the robustness of our downstream analyses while also providing tools for the initial investigation of the dataset. In the following sections 3.1.2 and 3.1.3, we present an overview of GLARE followed by the evaluation of its output in sections 3.2 and 3.3.

#### 3.1.2 Representation Learning

GLARE offers a range of widely applied representation learning techniques, including classic dimension reduction methods like PCA, t-SNE, and UMAP. However, GLARE also incorporates Sparse Autoencoder (SAE), a deep learning-based model that enables efficient data compression while preserving salient features. In this way, it can capture intricate hierarchical structures within the data by simultaneously learning both compressed data representation and the features necessary for reconstruction (Ng et al., 2011; Ranzato et al., 2007). The illustration of the overall pipeline and details of GLARE are shown in Figure 2.

**Figure 2.**
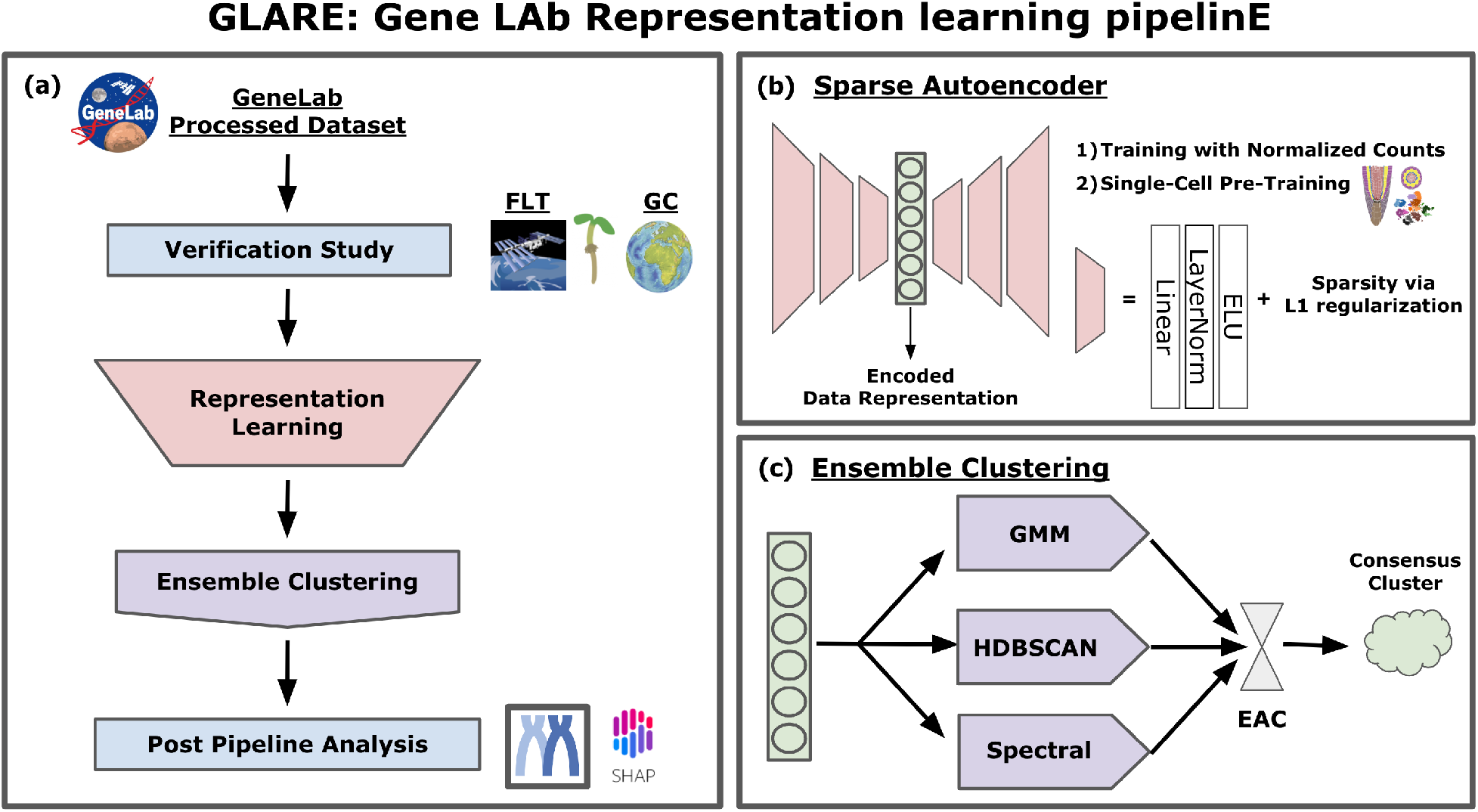
Overall pipeline of GLARE: Gene LAb Representation learning pipelinE. (a) Illustration of GLARE, starting with a verification study followed by representation learning and ensemble clustering. GLARE provides options for representation learning from PCA to state-of-the-art SAE pre-trained with high-throughput single-cell data. Retrieved data representation is then processed through ensemble clustering to find the hidden patterns within the data. Results from the verification study and ensemble clustering are then used for post-pipeline analysis. (b) Model architecture illustration of employed SAE for both training with and without pre-training. (c) Ensemble clustering using three base clustering algorithms based on different statistical methodologies. Evidence accumulation clustering is used to derive consensus clusters from these algorithms.

Our implementation of SAE is constructed with a sequence of building blocks, each comprising a Linear layer followed by LayerNorm and Exponential Linear Unit (ELU) activation. We chose to add the LayerNorm block to improve convergence and stable optimization, considering that our data consists of multiple experimental results from different environment settings (Ba et al., 2016). Towards this matter, we employ ELU activation as well (Clevert et al., 2015). We use three of these building blocks for the encoder and three blocks for the decoder to make the SAE. We choose L1-regularization over L2-regularization to induce the sparsity effectively and for better implicit feature selection (Ng, 2004), addressing the sparse and heterogeneous nature of normalized counts of FPKM values. The model training is optimized using mean squared error loss, Adam optimizer (Kingma and Ba, 2014) with weight decay, early stopping, and gradient clipping to address exploding gradients and ensure stable training. These hyperparameters were tested to find optimal parameter sets, which are described in our shared code repository (https://github.com/OpenScienceDataRepo/Plants_AWG/tree/main/Manuscript_Code/glare). Finally, after the model training, we extract data representation from the bottleneck layer between the encoder and decoder using this optimized model.

To further enhance the utility of these representations for downstream tasks such as clustering, we introduced an additional self-supervised learning step. We leveraged the pre-training step for the SAE with the addition of high-throughput single-cell data, specifically a single-cell root transcriptome dataset from Shulse et al. (2019), as the CARA dataset is drawn from root tip samples. This pre-training step complements the representations from the model by incorporating detailed single-cell transcriptome profiling of plant root cell types. We then take the pre-trained weights to fine-tune SAE using our normalized counts data to build Fine-Tuned SAE (FT-SAE). We maintain the original model structure and introduce adapter layers atop the main model to adjust varying dimensions between the single-cell matrix and our data appropriately, ensuring seamless integration into our SAE framework. We take the same procedure for the model optimization. The suggested self-supervised learning step offers several advantages, including the augmentation of feature granularity and the incorporation of cellular-level insights, thereby enhancing the fidelity and relevance of the learned representations for downstream analyses (Kiselev et al., 2018). Similar to our approach, such building of foundation models pre-trained with high-throughput single-cell data has demonstrated great utility in a diverse array of tasks in the life science field, including pattern recognition by incorporating foundational knowledge of the data (Hao et al., 2023).

#### 3.1.3 Ensemble Clustering

GLARE provides an ensemble clustering scheme to improve upon the commonly used application of single clustering approaches. Ensemble clustering offers several advantages over-relying on a single clustering algorithm. By merging the clustering outcomes from distinct statistical foundations through consensus voting, followed by hierarchical clustering with average linkage on the generated co-association matrix, we can mitigate the biases and noises inherent in each base clustering method to create more robust and reliable clustering results. Notably, when working with complex data such as representations retrieved from a fine-tuned sparse autoencoder, ensemble clustering can effectively address inherent complexities to capture hidden patterns and discover biologically meaningful clusters (Monti et al., 2003). GLARE adopts Evidence Accumulation Clustering (EAC) (Fred and Jain, 2005) as its ensemble clustering method, integrating three base clustering algorithms: GMM, HDBSCAN, and Spectral clustering. In addition to obtaining consensus cluster labels using EAC, researchers can leverage GLARE results from three base clustering algorithms to get unique clusters for each gene by retrieving the intersected cluster from its respective cluster assignments.

### 3.2 Data Representation Evaluation

In this section, we compare data representations from different algorithms that could be retrieved from GLARE. Figure 3, shows visualizations of each of the representations from FLT and GC using PCA, t-SNE, UMAP, SAE, and FT-SAE. Data representation from SAE and FT-SAE has n-dimensions depending on the number of neurons on the bottleneck layer. This value was determined through hyperparameter tuning, where various values of *n* were tested and evaluated based on their performance. The optimal value of *n* = 16 was selected based on the best reconstruction error observed during this tuning process. All other data representations from PCA, t-SNE, and UMAP have a 2-dimensional matrix. The PCA representations are highly condensed in a single region of the map, while t-SNE and UMAP representations exhibit a more widespread distribution. On the other hand, SAE and FT-SAE representations show more cluster-forming shapes for their t-SNE coordinates where the locally condensed points are separated from others.

**Figure 3.**
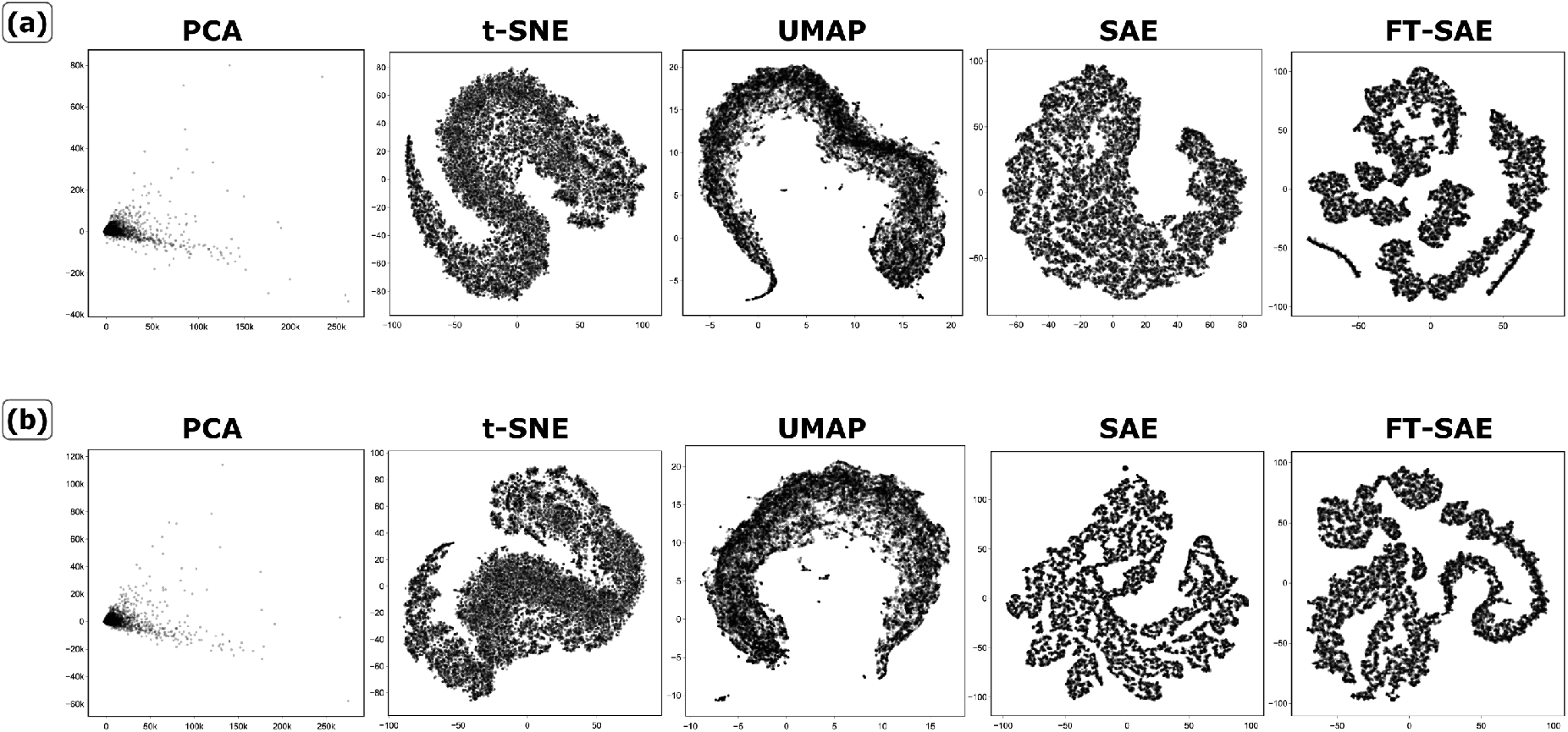
Comparison of data representations retrieved from GLARE. PCA, t-SNE, UMAP, SAE, and FT-SAE from left to right for both (a) FLT data (Upper-panel) and (b) GC data (Lower-panel). t-SNE was used for the visualization of 16-dimensional data representation from SAE and FT-SAE.

Although such an initial, qualitative visual analysis provides a useful overview of each technique’s output, GLARE provides for a more quantitative interpretation. Table 2 shows this next element of the analysis, evaluating these data representations using multiple quantitative measures: local neighborhood preservation through trustworthiness score (Venna and Kaski, 2001), retention of class-discriminative information through KNN classifier accuracy with cross-validation (Van Der Maaten et al., 2009), and quality of emergent clusters measured by the Silhouette Score (Rousseeuw, 1987) on k-means clustered data. Among the data representations that could be retrieved from GLARE, we compare all the methods that perform a non-linear transformation to the original dataset, leaving out PCA.

**Table 2:** Comparison of various evaluation metrics on data representations. FT-SAE shows the highest KNN accuracy and highest Silhouette score while having a lower trustworthiness score compared to others for both FLT and GC.

| Environment | Data Representations | Evaluation Metrics |  |  |
| --- | --- | --- | --- | --- |
| | | Trustworthiness Score ↑ | KNN Accuracy ↑ | Silhouette Score ↑ |
| FLT | t-SNE | 0.942 | 67.67 | 0.3842 |
|  | UMAP | 0.943 | 81.36 | 0.3985 |
|  | SAE | **0.967** | 90.66 | 0.4959 |
|  | FT-SAE | 0.888 | **96.08** | **0.5635** |
| GC | t-SNE | 0.946 | 64.04 | 0.3743 |
|  | UMAP | 0.949 | 80.14 | 0.3791 |
|  | SAE | **0.951** | 93.80 | 0.5634 |
|  | FT-SAE | 0.870 | **94.45** | **0.5886** |

Measuring the Silhouette score and performing the KNN classification on the labels from simple k-means clustering offers valuable insights into the quality of data representation, specifically, their utility in downstream tasks. Our results show that FT-SAE outperforms others on both of these measures, suggesting its robust capability to generate meaningful and discriminative representations. On the other hand, SAE shows the highest trustworthiness score for both FLT and GC. This result shows the advantages of nonlinear, sparse representation for complex biological datasets and its effectiveness in preserving local data structures. While FT-SAE retains a relatively high trustworthiness score with *>* 0.8 (Lee et al., 2007), it has the lowest among others. This is likely due to the model’s adaptation to features from the high-throughput single-cell dataset used in pretraining, which may not perfectly align with the local structure of the dataset used in the fine-tuning process. Specifically, the increased resolution of the single-cell data can induce clustering patterns that are not present in the CARA data, leading to discrepancies in how the model captures and represents the underlying structures of the dataset (Pan and Yang, 2009). However, this trade-off appears to be beneficial overall, with FT-SAE showing better performance in downstream tasks.

Overall, this analysis indicates FT-SAE is our method of choice for further analyses based on its KNN and Silhouette scores, which rank top of all techniques, and its acceptably high trustworthiness value. This suggests that the use of nonlinear, sparse representations from SAE, along with the introduction of the high-throughput single-cell dataset during pretraining, enhances the model’s ability to capture local and global structures in the data.

### 3.3 Clustering Results

Here, we present clustering results using GLARE on the CARA dataset. Figure 4 shows ensemble clustering results on the best-performing data representation, FT-SAE, with t-SNE used only for the purpose of visualization of the 16-dimensional data representation. We show individual clustering results from the base clustering algorithms we considered, GMM, HDBSCAN, and Spectral clustering, along with a final consensus cluster through evidence accumulation clustering (Fred and Jain, 2005). We note that GMM and spectral clustering require a user-defined cluster level.

**Figure 4.**
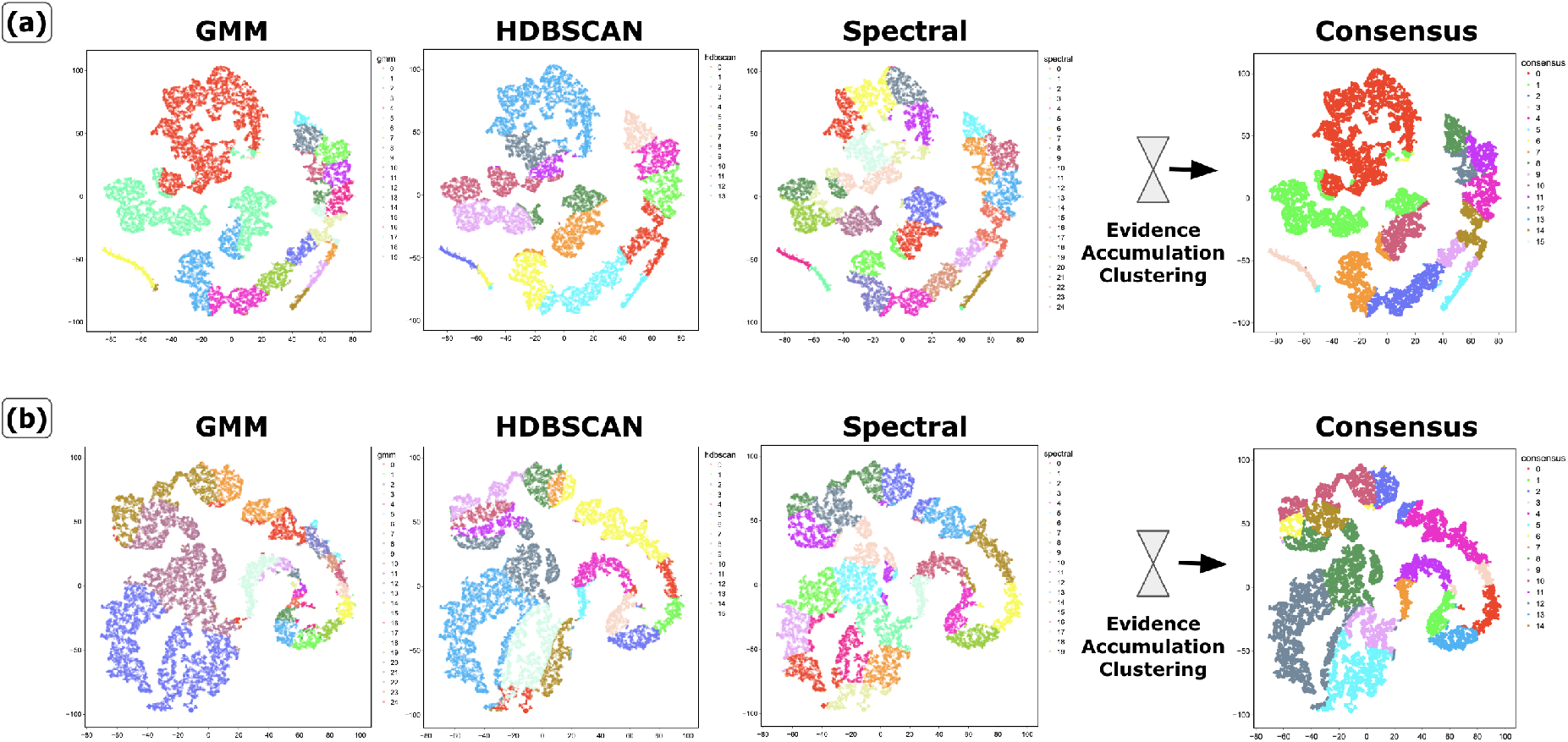
Ensemble clustering via EAC. Results from base clustering algorithms, GMM, HDBSCAN, and Spectral clustering, are shown starting from left to right for both (a) FLT FT-SAE representation (Upper-panel) and (b) GC FT-SAE representation (Lower-panel). t-SNE was used for the visualization of 16-dimensional data representation from FT-SAE. EAC results are shown at the right, with FLT having 16 consensus cluster labels and 15 consensus cluster labels for GC (depicted as different colors).

These were set to 20 and 25, respectively, for FLT and 25 and 20 for GC driven by results from previous studies (Shulse et al., 2019; Shahan et al., 2022). HDBSCAN defines its own cluster number.

#### Spaceflight

Clustering of the FLT dataset resulted in the identification of 20, 13, and 25 clusters for GMM, HDBSCAN, and Spectral clustering, respectively. GMM clusters had two large clusters, each containing 7,623 and 5,778 genes, with most of the other clusters having sizes of 300 to 1,000 genes. HDBSCAN showed a smaller number of clusters, where most of the clusters had 1000 to 2500 genes. Spectral clusters had the most consistent cluster sizes compared to GMM and HDBSCAN, with most of the clusters having 1,000 to 1,300 genes. These results highlight how the precise nature of clusters is different depending on the clustering approach taken. Each clustering strategy has distinct strengths. GMM works well when the data does not have well-defined boundaries, where HDBSCAN is useful for datasets with noise and outliers, and spectral clustering is highly suited for data with non-linear manifold structures (Xu and Tian, 2015). In order to leverage all of these advantages to a robust and reliable analysis of CARA data representation, we combined all three approaches via ensemble clustering through consensus voting (Vega-Pons and Ruiz-Shulcloper, 2011). These ensemble clusters exhibited diverse characteristics, having clusters with a size of *<* 1000 genes to two large clusters finding patterns in local structure, each containing 7,627 and 4,715 genes, similar to clusters identified by GMM. The number of clusters and size of remaining clusters, ranging from 1,000 to 2,500 genes, likely reflect the outputs of HDBSCAN and Spectral clusters, finding patterns throughout the global structure (Jain, 2010).

#### Ground Control

Clustering of the GC dataset resulted in the identification of 25, 15, and 20 clusters for GMM, HDBSCAN, and Spectral clustering, respectively. Despite the slight change in the number of clusters, the qualitative characteristics of the results remained largely consistent with those obtained from FLT. Specifically, GMM results revealed two large clusters, each containing 7,445 and 6,157 genes for GC, along with most of the other clusters having lesser sizes of 200 to 900 genes highlighting local patterns. Similarly, HDBSCAN and spectral clusters had a comparable consistent number of genes as FLT clusters, finding patterns throughout the global structure. Ensemble clustering demonstrated similar outcomes to FLT as well, exhibiting a diverse range of gene counts within each cluster.

### 3.4 Post Pipeline Analysis

Lastly, we demonstrate the full utility of GLARE using the results derived from the ensemble clustering on learned data representation of the CARA data and applying feature explanation analysis from the prediction task that we undertook for the verification study.

#### 3.4.1 Gene Ontology Analysis

Gene Ontology (GO) analysis, in conjunction with clustering results, is a widely used approach to find the functional significance of co-expressed genes in the clusters and provide a comprehensive understanding of the biological functions and processes underlying the observed gene expression patterns. We use the Metascape platform (http://metascape.org), which integrates various functional annotation databases (Zhou et al., 2019) to perform GO enrichment analysis. We take the clusters from EAC on FT-SAE and process them through Metascape after excluding clusters with extreme sizes, as this tool can only take gene lists of less than 3000 counts for the enrichment analysis. Specifically, we exclude the two large clusters for both FLT and GC datasets, along with one small cluster comprising only 2 genes in the FLT dataset, which leaves us 13 significant clusters for both FLT and GC. GO analysis on these clusters revealed various groups of ontologies, including cellular metabolic processes, oxidative phosphorylation, light response and signaling, and vesicle-mediated transport (Supplementary Table S1).

The prior study on the CARA dataset (Paul et al., 2017) found that genes associated with cell wall metabolism seemed most prevalent among the differentially expressed genes. Our analysis demonstrates that both FLT and GC clusters, which contain a high proportion of differentially expressed genes, are enriched in ontologies related to “Protein synthesis, Energy metabolism, and Stress response pathways” and “Cell wall biogenesis and Intracellular transport” as well. These results highlight pathways related to environmental adaptation and emphasize the critical role of cell wall metabolism identified in the CARA dataset. Moreover, we found unique hypoxia-related clusters that were only prevalent in FLT results. Root zone hypoxia is predicted to occur in spaceflight as a loss of buoyancy-driven convection in microgravity should limit oxygen resupply to intensely respiring tissues (e.g., Porterfield (2002)). However, transcriptional fingerprints of hypoxia response in plants in spaceflight have often proven elusive. We therefore concentrated the focus of the rest of our analysis on this hypoxic cluster. In Figure 5, we show a heatmap for the FPKM values for the genes within the main hypoxia cluster, GO analysis results for the hypoxia cluster using Metascape (Zhou et al., 2019) (Figure 5(b)), and Stress Knowledge Map (SKM) (Bleker et al., 2023) centered around the Transcription Factors (TFs) in the hypoxia cluster (Figure 5(c)). The Stress Knowledge Map (SKM; https://skm.nib.si/) is a curated resource offering two types of knowledge graphs on plant molecular interactions and stress signaling (Bleker et al., 2023). We used the Comprehensive Knowledge Network (CKN) to gain insights into stress signaling and associated plant biological processes around our genes of interest. The map in Figure 5(c) was drawn with five transcription factors (TFs) that we found in the 43 gene hypoxia cluster: *DREB2A, RHL41 / ZAT12, MYC2, RRTF1 / ERF109*, and *STZ / ZAT10*. The CKN map shows an intricate network of TFs and their interactions in the context of stress response mechanisms and related signaling pathways with other genes such as *HY5 / TED5, ABI1*, and *JAZ1*. Inspection of this network reveals ethylene as a likely important player in this response. GO analysis of the network for biological function (Supplementary Table S2) indicates elements of defense, water stress and cold response may also be important elements for further study.

**Figure 5.**
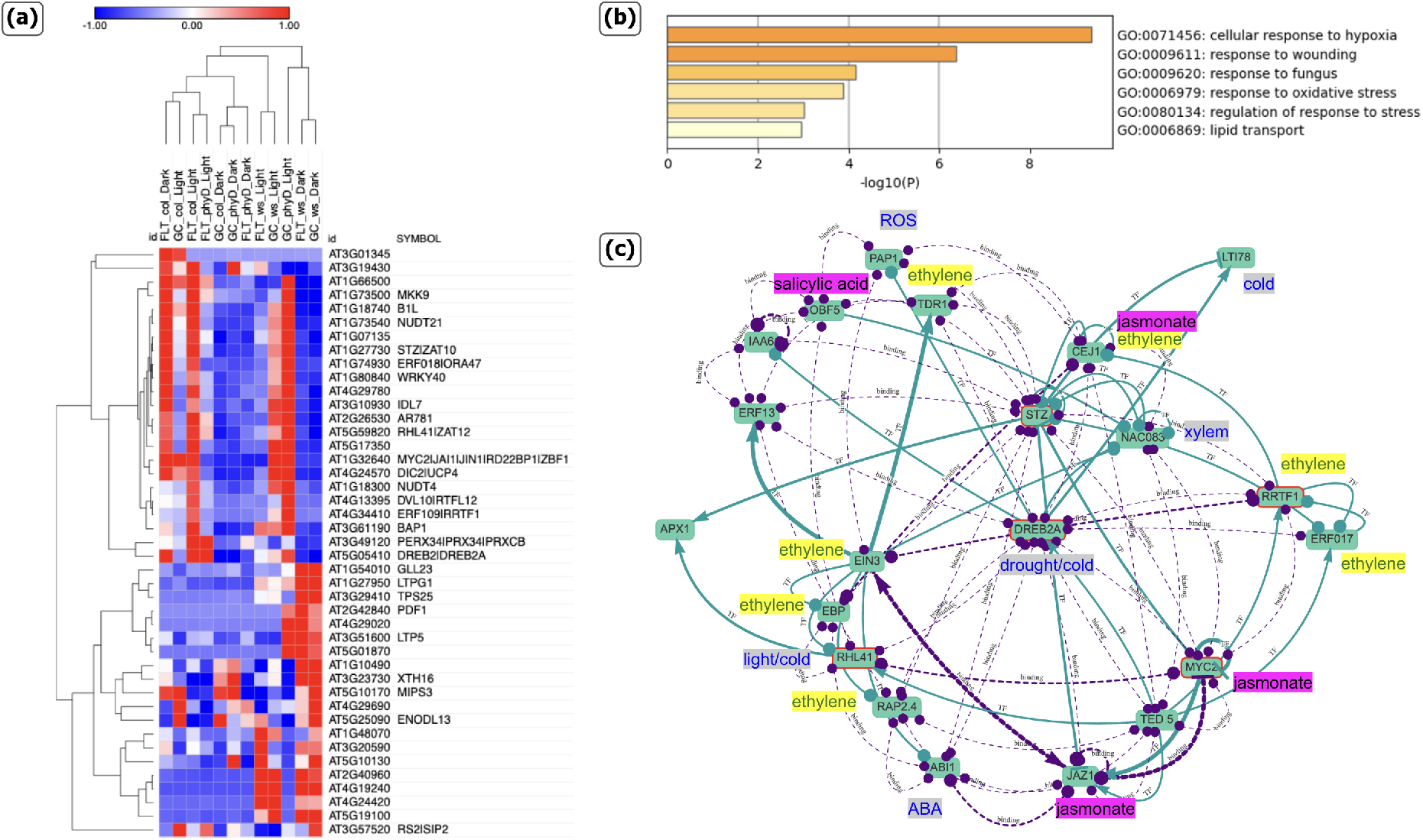
Analysis of hypoxia cluster found in FLT clustering result. (a) Heatmap of normalized FPKM values on hypoxia cluster. (b) Enriched ontology on hypoxia cluster from Metascape (c) Stress Knowledge Map (SKM) on five Transcription Factors (TFs) in hypoxia cluster: *‘DREB2A’, ‘RHL41 / ZAT12’, ‘MYC2’, ‘RRTF1 / ERF109’*, and *‘STZ / ZAT10’*. Purple lines: physical interaction (binding), teal lines: transcription factor interactions (TF). Annotations of relationships to Abscisic acid (ABA), Ethylene, Jasmonate, Light/cold, Reactive oxygen species (ROS), Salicylic acid and Xylem are from manual inspection of each gene’s annotated roles in The Arabidopsis Information Resource (https://www.arabidopsis.org).

Additionally, we identified that the hypoxia cluster had the highest percentage of differentially expressed genes within the cluster (DEG proportions in Supplementary Table S3). These genes were then used for cell-type prediction using scPlantDB (He et al., 2024) to find highlighted cells in different root organ groups, taking advantage of the single-cell pre-trained model FT-SAE of GLARE. We perform this cell-type prediction for all the clusters as well (Supplementary Table S3). The hypoxia cluster had more than half of the differentially expressed genes identified in the lateral root cap and columella root cap, while also appearing in phloem parenchyma and xylem. These results reveal a distinct enrichment of hypoxia-responsive genes in root cap tissues, while also highlighting significant involvement of the root vasculature, suggesting a coordinated response to low oxygen conditions across root structures.

#### 3.4.2 SHAP Analysis

Up to this point, our analysis has been directed toward uncovering distinct patterns, specifically between FLT and GC, by generating separate data representations for clustering and GO analysis. However, we chose CARA as a dataset to interrogate due to the multiple experimental factors within the experiment’s design. Therefore, after identifying the patterns within the FLT data using GLARE, particularly a hypoxia cluster, we used this newly identified cluster to evaluate the effect of varying light conditions on different genotypes in each location. We took the found TFs within the hypoxia cluster and applied SHAP analysis to quantify feature contribution, thereby explaining which experimental conditions had the most effect in classifying this pattern within the data between FLT and GC. SHAP analysis provides a way to understand the impact of each feature on the model’s predictions, enabling better model transparency and insights into the underlying relationships within the data (Lundberg and Lee, 2017). Higher positive SHAP scores reflect features contributing more to this discrimination within the dataset to designate a sample to FLT, while negative values reveal factors that have a negative impact on the FLT assignment, i.e., reveal the data as GC.

In Figure 6(e), we show the difference in SHAP values for the TFs within the hypoxia cluster. Among these five TFs, *ZAT12* has the largest aggregate difference in SHAP values between FLT and GC, and *MYC2* the smallest. In Figure 6(a) to (d), we show SHAP local bar plots for *ZAT12* and *MYC2* explaining the feature importance representing different environmental conditions and genotypes. For both *ZAT12* and *MYC2*, the *PHYD* mutants and Col-0 genotype in the dark setting had the most contribution to model prediction in FLT (Figure 6(a) and (c)). On the other hand, both the dark and light settings for the Col-0 genotype had the most contribution to model prediction in GC (Figure 6(b) and (d)). The large difference in aggregated SHAP value between FLT and GC for *ZAT12* suggests that the relative importance and contributions of these features vary significantly between the FLT and GC. In contrast, the contributions for *MYC2* appear more consistent and stable across both FLT and GC classifications. This analysis using SHAP values provides deeper insight into the differentiation between environmental conditions and genotypes at different locations, solely based on gene expression data. Lastly, in Figure 7, we present summary SHAP plots on these features, varying light conditions on different genotypes, to offer a more comprehensive understanding of feature contribution across the entire dataset.

**Figure 6.**
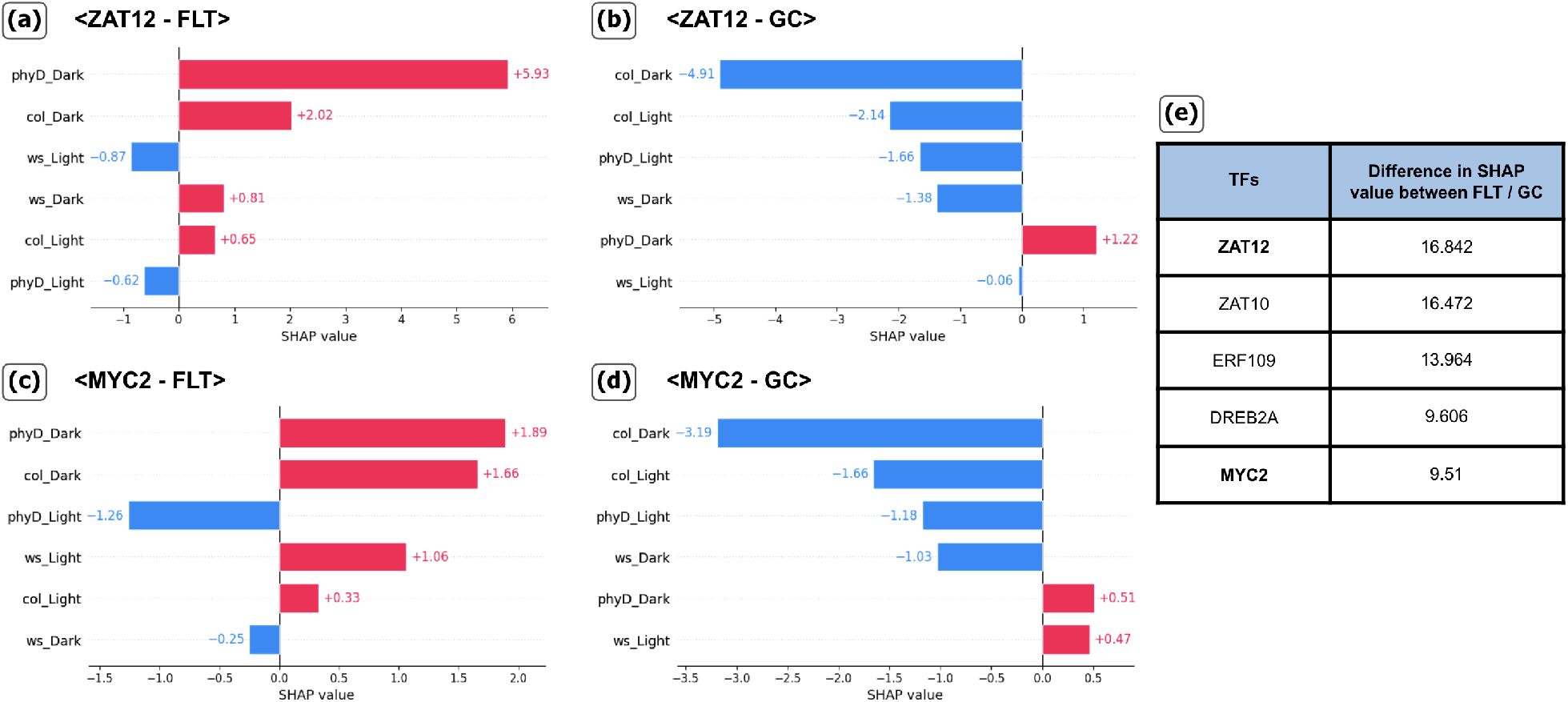
SHAP analysis on Transcription Factors (TFs) in the hypoxia cluster. A positive SHAP value (Red color) means that the feature value made a greater contribution than others in classifying the gene as FLT, while a negative SHAP value (Blue color) suggests they had more contribution in GC classification. (a) *ZAT12* - FLT (b) *ZAT12* - GC (c) *MYC2* - FLT (d) *MYC2* - GC (e) Summary of difference in SHAP value between FLT and GC for the 5 TFs in hypoxia.

**Figure 7.**
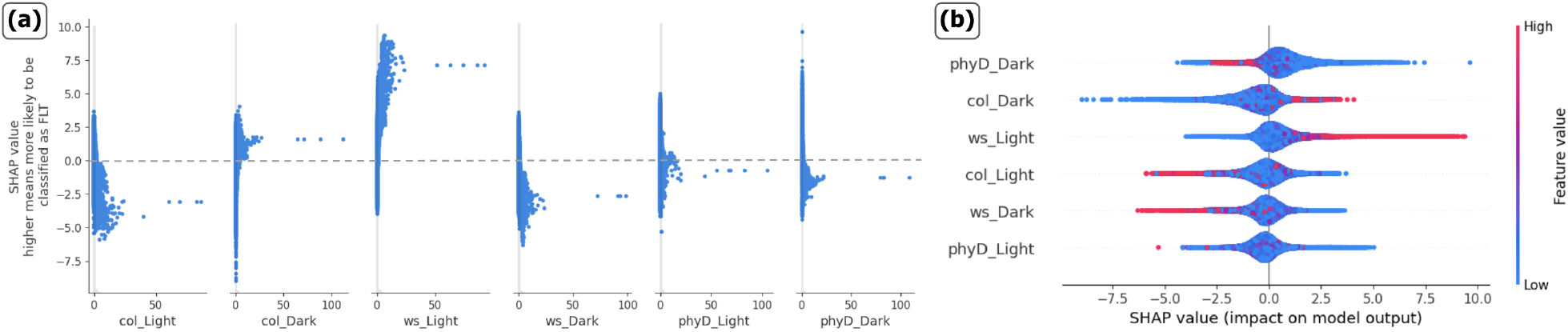
SHAP value distribution for each treatment. Comparing SHAP values from a classification using the XGBoost on the restructured CARA dataset. (a) The summary SHAP value scatterplot for each feature displays the distribution of SHAP values alongside raw feature values. (b) The summary SHAP beeswarm plots, where features are ordered by their importance (measured by mean absolute SHAP values), with the most impactful features appearing at the top. The color bar represents raw feature values. Both plots present the same information.

We can observe features with different degrees of impact on the model’s prediction from the SHAP value scatterplot in Figure 7(a), for example, the WS genotype: in a light setting, the majority of the data aligns with positive SHAP values, supporting FLT classification, whereas under dark conditions, the trend is reversed. The beeswarm plot (Figure 7(b)) illustrates the distribution of SHAP values for each feature as well. The color gradient from blue to red represents the feature value (FPKM values), with blue indicating low expression and red indicating high expression. Figure 7(b) illustrates that *PHYD* mutants in a dark setting have the highest effect on the classification with longer tails towards positive SHAP value, while most of the high FPKM values have negative SHAP value. Suggesting that high expression levels from *PHYD* mutants in dark settings decrease the likelihood of FLT classification. Similarly, the Col genotype in a dark setting has tails toward negative SHAP values, while most of the high FPKM values have positive SHAP values. These observations emphasize the presence of intricate interactions between gene expressions, reflecting the complexity of the transcriptome data and the underlying biological mechanisms.

Such SHAP analysis offers a unique perspective on the patterns within the dataset, particularly when the data comprises various environmental settings as features. Through analyzing the variations and similarities in SHAP values, researchers can identify genes that are sensitive to complex environments based on genotype-dependent patterns in the data.

## 4 Discussion

In this study, we present an analysis pipeline, GLARE, that employs a state-of-the-art representation learning model with self-supervised learning. We chose a previously analyzed dataset, the CARA experiment (OSD-120), which allows for an investigation of the overall utility of the pipeline itself and a comparison with the prior findings. For analysis of the root samples in the CARA spaceflight data, we pre-trained the model using high-throughput plant root single-cell data, along with ensemble clustering, to identify hidden patterns in the spaceflight transcriptome. For other spaceflight datasets, such as whole seedlings, shoot tissues, microbe, animal tissues, or cell types, matching pre-training datasets to the particular experimental design would similarly add significant depth to these analyses. After the full pipeline, we present a recommended addition for post-pipeline analysis employing select bioinformatics tools and adding post hoc explainability to the deep learning approach by applying SHAP analysis. Such analyses confirmed previous patterns found in the data, such as cell wall remodeling and vesicle-mediated transport, but critically revealed new features, notably a molecular signature of hypoxic stress in the spaceflight samples that is predicted from the lack of buoyancy-driven convection in spaceflight but that has proven complex to extract from many plant transcriptomic datasets. However, our analyses also revealed that this cryptic signature was dependent on experimental conditions such as plant genotype and lighting regime. For example, Figure 6 shows that SHAP analysis of the 5 signature spaceflight-related, hypoxia-response transcription factors identified in this study can potentially help explain the complex interplay of genotype and lighting conditions that make these signals complex to identify in current spaceflight datasets without machine learning interrogation.

Although we present one post hoc analysis pipeline for the output of GLARE, researchers can readily leverage their preferred analytics tools when applying GLARE to their datasets to uncover patterns. To this end, we actively encourage contributions and novel suggestions through our open science repository. Its open-source nature means researchers can readily adapt GLARE on other datasets from GeneLab and elsewhere to reinforce their initial studies and expand on these computational findings. The recent rapid advancement in the machine learning field warrants future work on GLARE. Similar to our approach, integrating single-cell datasets has begun to be widely adopted for their advantage in providing nuanced insights to the cellular level. Indeed, transformer-based foundation models for single-cell multi-omics have been suggested (Cui et al., 2024), which offer the potential to generate synthetic data or for gene network inference. Our future vision for GLARE is to extend beyond autoencoder-based models to add more advanced self-supervised representation learning models, such as the contrastive learning methods that are well-used in the field of computer vision and natural language processing (Chen et al., 2020) to enhance robustness for smaller datasets with fewer features. Additionally, causal representation learning methods offer an exciting avenue to discover the causal relationship between related genes (Uelwer et al., 2023; Schölkopf et al., 2021).

## Author Contributions

DH.S. and R.B. conceived of the study and fundamental design. DH.S. and H.F.S. contributed to model testing, data analysis, and figure preparation. DH.S., H.F.S., M.Z., R.B., A.-L.P, R.J.F., and S.G. contributed to manuscript preparation. All authors contributed to the manuscript review and editing.

## Acknowledgments

The authors would like to acknowledge the sequencing and bioinformatics services provided by the Interdisciplinary Center for Biotechnology Research’s (ICBR) Gene Expression (RRID:SCR_019145), NextGen Sequencing (RRID:SCR_019152), and Bioinformatics (RRID:SCR_019120) cores.

## Data Availability Statement

The dataset (OSD-120) utilized in this method can be found on the NASA GeneLab Data System (https://genelab.nasa.gov/). The code utilized for data analysis can be found on the publicly available GitHub repository (https://github.com/OpenScienceDataRepo/Plants_AWG/tree/main/Manuscript_Code/glare).

## Supplementary Data and Table

We provide supplementary data at the OSDR GitHub repository (https://github.com/OpenScienceDataRepo/Plants_AWG/tree/main/Manuscript_Code/glare), including the codes for the method and results such as single-cell pre-trained model weights, data representations, ensemble clustering results, Gene Ontology (GO) analysis results for all clusters, and predicted SHAP values for both FLT and GC. All the Supplementary Tables demonstrating further details of our results are also included in the repository.

### Preprocessing

GLARE provides tools for the initial investigation of the dataset by employing the conventional dimensionality reduction approach of PCA to achieve simple data representation followed by k-means clustering algorithm. Figure S1, shows the distribution of the principal components and clustering results on them for the CARA data. Notably, results of both spaceflight (FLT) and ground control (GC) experiments exhibit similar distributions and clustering patterns, characterized by a concentration of data points within a single cluster. We took this opportunity to perform cluster-based outlier detection (Sikder and Batarseh, 2023), discarding out-of-distribution clusters and keeping the concentrated clusters. For both FLT and GC data, we take the three most concentrated clusters, discarding only 3 data points, to the representation learning step of the pipeline.

**Figure S1:**
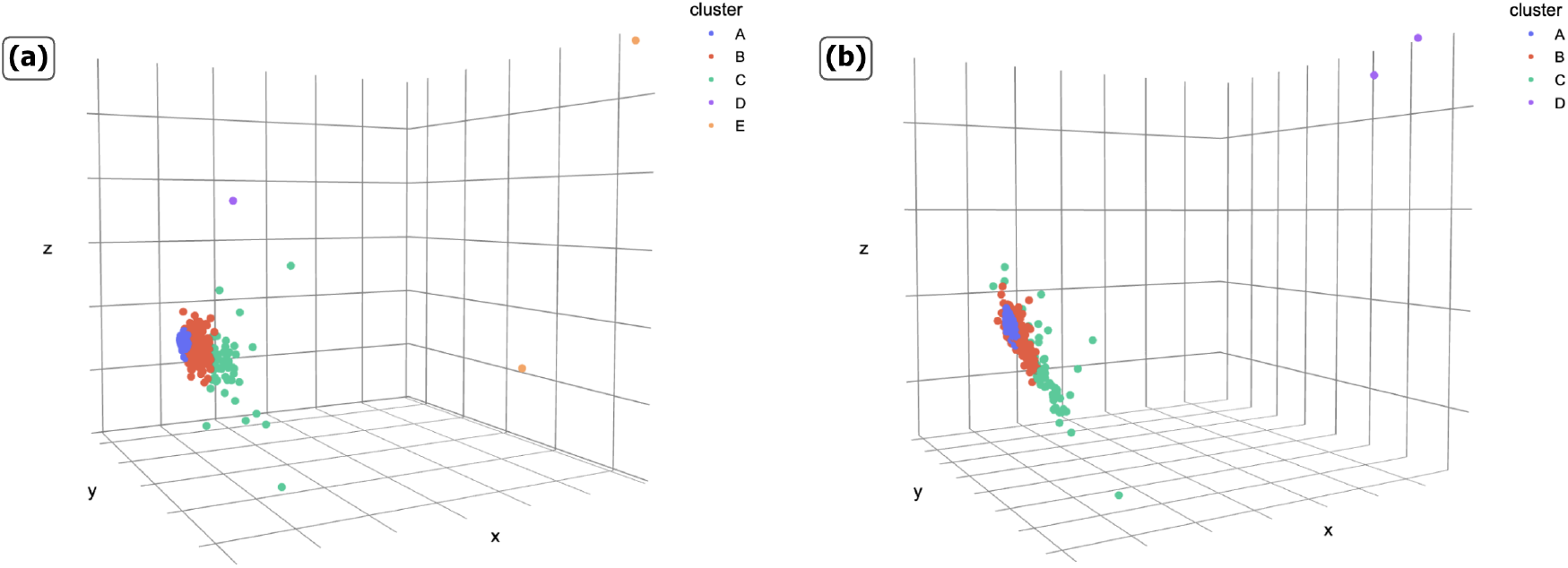
Initial investigation via PCA and k-means. (a) Clustering result on spaceflight (FLT) data (without any data melting) PCA (*n*_*components* = 3). (b) Clustering results on ground control (GC) data (without any data melting) PCA (*n*_*components* = 3). ∼ 98% of the data is clustered on cluster *A* (blue) for both FLT and GC. In this study, from the *>*25,000 genes in the datasets, we only discard three genes (cluster D and E in (a) and cluster D in (b)) for both FLT and GC that are separated from concentrated clusters A, B, and C as a means of outlier detection. These genes are: AT1G0759, AT3G41768, ATMG00020.

While our study employed cluster-based outlier detection, the applicability of this approach may vary across different GeneLab datasets. To address this variability, we have integrated additional outlier detection methods that researchers can employ using GLARE, including z-score analysis on the first principal component and the Local Outlier Factor (LOF) method (Jolliffe and Cadima, 2016; Breunig et al., 2000). Researchers can utilize PCA and k-means clustering for preliminary investigation, as well as to detect outliers as we demonstrated with CARA dataset, and then apply suitable outlier detection techniques tailored to their specific GeneLab datasets.

